# Decoding semantic predictions from EEG prior to word onset

**DOI:** 10.1101/393066

**Authors:** Edvard Heikel, Jona Sassenhagen, Christian J. Fiebach

## Abstract

**ABSTRACT:** The outstanding speed of language comprehension necessitates a highly efficient implementation of cognitive-linguistic processes. The domain-general theory of Predictive Coding suggests that our brain solves this problem by continuously forming linguistic predictions about expected upcoming input. The neurophysiological implementation of these predictive linguistic processes, however, is not yet understood. Here, we use EEG (human participants, both sexes) to investigate the existence and nature of online-generated, category-level semantic representations during sentence processing. We conducted two experiments in which some nouns – embedded in a predictive spoken sentence context – were unexpectedly delayed by 1 second. Target nouns were either abstract/concrete (Experiment 1) or animate/inanimate (Experiment 2). We hypothesized that if neural prediction error signals following (temporary) omissions carry specific information about the stimulus, the semantic category of the upcoming target word is encoded in brain activity prior to its presentation. Using time-generalized multivariate pattern analysis, we demonstrate significant decoding of word category from silent periods directly preceding the target word, in both experiments. This provides direct evidence for predictive coding during sentence processing, i.e., that information about a word can be encoded in brain activity before it is perceived. While the same semantic contrast could also be decoded from EEG activity elicited by isolated words (Experiment 1), the identified neural patterns did not generalize to pre-stimulus delay period activity in sentences. Our results not only indicate that the brain processes language predictively, but also demonstrate the nature and sentence-specificity of category-level semantic predictions preactivated during sentence comprehension.

**STATEMENT OF SIGNIFICANCE:** The speed of language comprehension necessitates a highly efficient implementation of cognitive-linguistic processes. Predictive processing has been suggested as a solution to this problem, but the underlying neural mechanisms and linguistic content of such predictions are only poorly understood. Inspired by Predictive Coding theory, we investigate whether the meaning of expected, but not-yet heard words can be decoded from brain activity. Using EEG, we can predict if a word is, e.g., abstract (as opposed to concrete), or animate (vs. inanimate), from brain signals preceding the word itself. This strengthens predictive coding theory as a likely candidate for the principled neural mechanisms underlying online processing of language and indicates that predictive processing applies to highly abstract categories like semantics.

## INTRODUCTION

Predictive coding has become an influential theory of neural information processing. It postulates that the brain continuously predicts incoming sensory information based on generative models of their (hidden) sources (e.g., Friston, 2010; Clark, 2013). Predictive coding assumes that ‘ explaining away’ predictable input minimizes information-processing requirements. As a consequence, new sensory input elicits prediction error signals representing the difference between predicted and actual input. According to this framework, these prediction errors are passed on to hierarchically higher brain areas in order to update the internal generative model, thereby optimizing future predictions.

The remarkable speed of language processing implies the possibility of predictive mechanisms also in sentence comprehension (see Kuperberg and Jaeger, 2015; Lewis and Bastiaansen, 2015, for recent discussions). Many established neural correlates of language processing seem to be plausibly associated with the concept of a prediction error (such as the N400 event-related brain potential/ERP typically elicited by violations of context-dependent semantic expectations; cf. Kutas et al., 2011; Rabovsky and McRae, 2014; Frank et al., 2015). Research in psycho- and neurolinguistics has focused on whether or not higher-level (lexical, semantic) representations can predictively pre-activate lower-level (phonemic or orthographic) features to facilitate language processing (van Petten and Luka, 2012; Kuperberg and Jaeger, 2015). Neural markers of prediction violations have been interpreted indicating pre-activation of, e.g., specific word forms (DeLong et al., 2005; but see Nieuwland et al., 2017) or lexical/semantic word features (Szewczyk and Schriefers, 2013; Wicha et al., 2004; Kwon et al., 2017). However, looking closely, this research paradigm only shows *that* the brain predicts during language processing – not *what* it predicts (Ito et al., 2017). Mechanistic models of language, however, require a more direct understanding of whether and how linguistic predictions are neurally encoded during sentence processing – i.e., directly probing the content of predictions in rich linguistic contexts.

As noted above, any sensory input will induce a prediction error signal coding its deviation from current internal predictions. If an expected stimulus is entirely absent, the prediction error should, in principle, only contain information about the prediction (Kok and De Lange, 2015). This provides a direct window for examining the nature of internal predictions, e.g., by occluding (Smith and Muckli, 2010) or omitting (SanMiguel et al., 2013) sensory information of a target stimulus. Applying this logic to language, Bendixen et al. (2014) demonstrated specific auditory cortex responses to omissions of word segments. Bonhage et al. (2015) and Boylan et al. (2014) used functional MRI (fMRI) to explore brain activation patterns associated with category-level (noun/verb) predictions of target words delayed for several seconds. However, due to the low temporal resolution of fMRI and the artificial nature of these visual delayed-target paradigms, these results contribute little to understanding predictive online-sentence processing. To the best of our knowledge, the existence of semantic-level predictions during online sentence processing and their underlying neural implementation have so far not been directly investigated using this approach.

We propose here that if specific semantic features of words are predictively pre-activated during sentence comprehension, they should be decodable from brain activity during the period of silence as an expected word is briefly and unexpectedly delayed – because as derived above, the prediction error signal elicited by this (temporary) omission should carry information about the word’s meaning. To test this, we combined a temporally precise technique, EEG, with multivariate pattern analysis (MVPA). First, we aimed at decoding target noun concreteness, because concreteness has a robust neural signature (Dufau et al., 2015; Huang, Lee, & Federmeier, 2010; West and Holcomb, 2000). We additionally examined whether the predictive neural encoding of semantic category observed in a sentence context is comparable to the semantic representation elicited when target nouns are presented in isolation. We then replicated in a second, independent experiment the decoding of semantic category from pre-target pauses, with noun animacy as the to-be-decoded semantic feature.

## METHODS EXPERIMENT 1: DECODING WORD CONCRETNESS FROM SILENCE-EVOKED EEG

Analysis scripts used for this report are accessible under https://github.com/heikele/sempred. Data will be uploaded to Zenodo no later than time of publication.

### Participants

Forty-five native speakers of German were recruited, who reported no history of neurological or psychiatric disorders and provided informed consent according to protocols approved by the local ethics committee prior to participating. Five participants were excluded from further analysis due to faulty equipment or software error (a broken ground electrode or an incorrect trigger signal). Two participants were excluded because the number of rejected trials during artefact correction exceeded the threshold of 33% (i.e. < 100 remaining trials). Furthermore, one non-native speaker was excluded and two further participants because not all three experiments (reported below) could be completed. The final sample consisted of *n* = 35 participants (mean age = 22.02, SD = 4.87; age range 18 to 42; 6 males).

### Experimental Design: Stimuli and procedure

Participants were seated in front of a monitor in a semi lit room and listened to sentences played electronically at a comfortable loudness. Two speakers were placed on either side of the screen. Trials started with a fixation cross and, after a 400-600 ms delay, were followed by the auditory sentence presentation. Sentence length varied between 2.59 and 6.38 s (mean = 4.06 s, SD = .87). To ensure semantic-level processing, participants judged directly after each sentence via button press whether or not it was synonymous with the immediately preceding sentence (space bar: synonymous, any other key: non-synonymous). A warning tone was played when a wrong answer was given.

In total, 350 sentences were presented, of which 150 contained a pause of 1 s directly preceding a target word. Of these critical sentences, 75 constrained for an abstract, and 75 for a concrete target noun (see Table 1 for examples). The sentences varied between 4 and 16 words in length (abstract: range 5 to 16, mean = 8.97, SD = 2.82; concrete: range 4 to 16, mean = 9.21, SD = 2.52), with the delayed word occurring 97 times in the final (abstract: 45, concrete: 52), 28 in the penultimate (abstract: 14, concrete: 14), and 25 times in an earlier position (abstract: 16, concrete: 9). 160 further sentences served as filler items. 40 sentences were additionally included as targets for the synonym task. Concrete words were about as frequent as abstract words (log counts per billion, using the German SUBTLEX-DE, Brysbaert et al., 2011: concrete, mean = 5.74, SD = 1.54. abstract, mean = 5.98, SD = 1.81). Mean target word cloze probability was moderate to high (mean = .57; SD = .36; cf. Block and Baldwin, 2010), according to ratings provided by an independent sample (n = 15; note that concrete words had a somewhat higher cloze rating than abstract words on average; concrete: mean = .66, SD = .35; abstract: mean = .47, SD = .35). Importantly, constraint for the to-be-decoded semantic target category (i.e., the proportion of abstract words given if the target word was abstract, or concrete words if the target word was concrete) was very high, and similar across the two conditions (abstract: mean = .94, SD = .17, concrete: mean = .99, SD = .05, respectively).

**Table 1.** Examples of critical sentences used in each experiment, with inserted pauses indicated by square brackets.

| Condition | Example Sentence |
| --- | --- |
| <u>Experiment 1</u> |  |
| Abstract | In einem Leserbrief bringt sie ihre [...] **Meinung** zum Ausdruck.<br>*In a letter to the editor, she expresses her opinion.* |
| Concrete | Sie stellte den Teller auf dem Tisch ab und nahm Messer und [...] **Gabel** in die Hand.<br>*She set the plate down on the table and picked up the knife and fork.* |
| <u>Experiment 2</u> |  |
| Animate | Um sich auszuheulen, ruft Anne ihre <u>beste</u> [...] **Freundin** an.<br>*To herself cry-out, calls Anne her <u>best</u> [...] **friend**<sub>female</sub> on.*<br>*To talk it out and get it off her chest, Ann calls her best friend.* |
| Inanimate | Weil die Sonne sie blendet, setzt Kira ihre <u>beste</u> [...] **Sonnenbrille** auf.<br>*Because the sun her blinds, puts Kira her <u>best</u> [...] **sunglasses** on.*<br>*Because the sun blinds her, Kira puts on her best sunglasses.* |

Bold print indicates the target words for which cloze probability was determined (see text).
Square brackets indicate the positions preceding these target nouns, where a delay of 1,000 ms was inserted, and underlined words denote the identical pre-target words preceding the delay in Experiment 1. Literal translations are printed in italics, and word-for-word translations (indicating the identical context word preceding the pause in Experiment 2) are given in courier font.

In the same session as the sentence experiment, two additional experimental runs were acquired. In these, participants were presented with isolated spoken (Experiment 2B) and printed (Experiment 2C) words (150 words per list; 75 per condition), to examine the sensitivity of our analysis pipelines for detecting imageability effects outside of sentence context and to test cross-decoding between the different experiments. None of the stimuli in these experiments appeared in the sentence study, and each word was presented only once. The words were presented in random order while participants performed a 1-back synonymity judgment task (analogous to the sentence experiment) – i.e., indicated for each word whether or not it was synonymous with the previous word. Twelve additional target words for the judgment task were presented in each experiment; these items were excluded from all analyses. The word lists were matched pairwise between conditions, within each experiment, by the number of letters (auditory: mean = 5.76, SD = 1.34 for both abstract and concrete words; visual: mean = 5.44, SD = 1.24 and mean = 5.44, SD = 1.25 for abstract and concrete words, respectively) and word frequency (auditory: mean = 6.84, SD = 1.25 and 6.86, SD = 1.23; visual: mean = 6.99, SD = 0.79 and 6.99, SD = 0.78 for abstract and concrete words, respectively). Visual word lists were additionally matched on orthographical neighbourhood density (via the OLD20 measure, see Yarkoni, Balota, & Yap, 2008). OLD20 was calculated by averaging the Levenshtein distance between a word and its 20 closest orthographic neighbours, using the SUBTLEX-DE (Brysbaert et al, 2011), for both abstract (mean = 1.70, SD = 0.43) and concrete (mean = 1.67, SD = 0.44) visual target nouns. Auditory words were also matched by spoken word length (abstract: mean = 644 ms, SD = 95; concrete: mean = 561 ms, SD = 79).

### EEG Data Acquisition and Pre-Processing

Electroencephalographic (EEG) data were collected using 64 active Ag/AgCl electrodes (ActiCAP; Brain Products GmbH, Gilching, Germany) arranged in an extended 10-20 layout, using either a brainAmp or an actiChamp amplifier (both: Brain Products GmbH, Gilching, Germany); sampling rate of 1,000 Hz, low-pass filter of 500 Hz, time constant of 10s, and left earlobe as ground. Off-line, data were re-referenced to the linked mastoids. Pre-processing and all further analyses were conducted with MNE-Python (Gramfort et al, 2014) and MVPA pipelines were implemented using MNE-Python in conjunction with scikit-learn (Pedregosa et al, 2011). To speed up the computationally intensive multivariate analyses, data were down-sampled to 200 Hz. Pre-processing involved a high-pass filter of .1 Hz and a low-pass filter of 30 Hz, segmentation into epochs from -.3 to 1 s relative to the start of the critical event (i.e., the pause onset in the sentences and the word onset in the single word runs), and baseline-correction (-.3 to 0 s). Independent component analysis (Jung et al, 2000) was used to correct eye-movements and muscle artefacts. An automated trial rejection/repair method (Jas et al, 2017) was applied. Then, peri-ocular (SO1, SO2) and mastoid electrodes were discarded, leaving 60 electrodes for all further analyses.

### Statistical Analysis: Event-related Brain Potentials

Event-related brain potentials (ERPs) were analysed with a non-parametric cluster-level permutation test across subjects (Maris and Oostenveld, 2007). Subject-specific averages were calculated for each condition, and difference waves created (i.e., concrete minus abstract). Significant spatio-temporal clusters were identified in the empirical data. Then, across 1,024 permutations, surrogate values under the null hypothesis were created by random sign flipping. Observed clusters exceeding the 2.5th and 97.5th percentiles in the surrogate data were considered to significantly differ from zero at the 5% level. Time points in the pre-stimulus baseline were excluded from statistical analysis in order to maximize power (Groppe et al., 2011).

### Statistical Analysis: Time-Generalized Multivariate Pattern Analysis

Multivariate pattern analysis (MVPA) was used to test whether the semantic category of the upcoming word – which was constrained for by the preceding sentence context, see above – could be predicted (henceforth: decoded) from brain activity elicited during the pre-target pause. As we had few specific a-priori expectations concerning the temporal extent of the hypothesized prediction error signal, time-resolved MVPA was conducted using generalization across time (GAT) decoding as outlined by King and Dehaene (2014; see also Heikel et al., 2018, for a previous application of this procedure in the linguistic domain from our group). Specifically, for each subject and at each of the 261 time points in the epoch (i.e., 1,300 ms sampled at 200 Hz), a linear Support Vector Machine classifier (SVM, with default penalty parameter: C=1.0) was trained to classify the semantic category of the target noun (abstract vs. concrete) based on the EEG pattern elicited during the pause preceding the target noun.

To ensure statistical independence, a 5-fold stratified cross-validation procedure was used. Specifically, over five iterations, the classifier was trained on 4/5ths of the data and tested on the remaining fifth fold, so that each fold was used for testing once. On the testing folds, the classifier was not only tested on the time point it was trained on, but also on all other time points in the epoch, thereby assessing whether it generalizes across time.

Classifier performance was scored by calculating the Area under the Curve of the Receiver-Operating Characteristic (ROC-AUC; Hastie et al., 2009), and then averaged across folds. The resulting score corresponds to the rate of true relative to false positives, weighted by the confidence of this classification, per combination of training and testing time points. More specifically, here, they correspond to the degree to which the EEG signal at each time point in the delay period contains information about the semantic category of the delayed word; above-chance decoding at one time-point indicates that the categorical concreteness of the delayed word (i.e., concrete vs. abstract) can indeed be read out from brain activity at this time point.

These decoding accuracy scores can be visualized in a subject-specific generalization-across time (GAT) matrix with training time on the y-axis and testing time on the x-axis. The GAT matrix can provide information about whether a neural pattern elicited at a certain time point in the trial is temporally restricted to the training time, or extends to other time points, or possibly even re-occurs at later time points (see King and Dehaene, 2014 for details; see also Heikel et al., 2018). Decoding scores (excluding, analogous to the ERP analysis, baseline data) were statistically evaluated at the group level by testing against chance (.5) at all time points using Threshold Free Cluster Enhancement (TFCE, h = .2, e = .1; Smith and Nichols, 2009). The same MVPA pipeline was used for the words presented in isolation (visual and auditory).

Additionally, in order to understand better the nature of predictive neural representations associated with the expectation of concrete vs. abstract words, classifiers were not only tested within the modality that they were trained in. We also examined whether classifiers trained to decode semantic category under conditions of single word presentation, can decode the concreteness category from the EEG activity measured during the pre-word pause in the sentence experiment.

## RESULTS EXPERIMENT 1

### Decoding Concrete- vs. Abstract Nouns prior to their Presentation

ERPs elicited during pauses preceding expected concrete vs. expected abstract nouns did not differ (Figure 1A & B). Accordingly, cluster-level permutation testing revealed no significant ERP differences (Figure 1A). However, using time-generalized MVPA we could reliably decode the semantic category status, i.e., abstract vs. concrete, of the delayed target word, from the pause interval preceding its presentation. Along the diagonal, decoding exceeded chance to a significant degree from 280 to 610 ms after the onset of the pause, as identified by threshold free cluster enhancement (TCFE, p < .05; Figure 1C for the GAT matrix and clusters of significance). Time-generalized decoding, in addition, indicated a temporally extensive, roughly square pattern of decoding. I.e., decoders trained on different time points could successfully predict concreteness condition relatively broadly between 200 and 1000 ms. This indicates that one and the same neural pattern carrying information about the semantic difference persisted for an extended time span.

**Figure 1.**
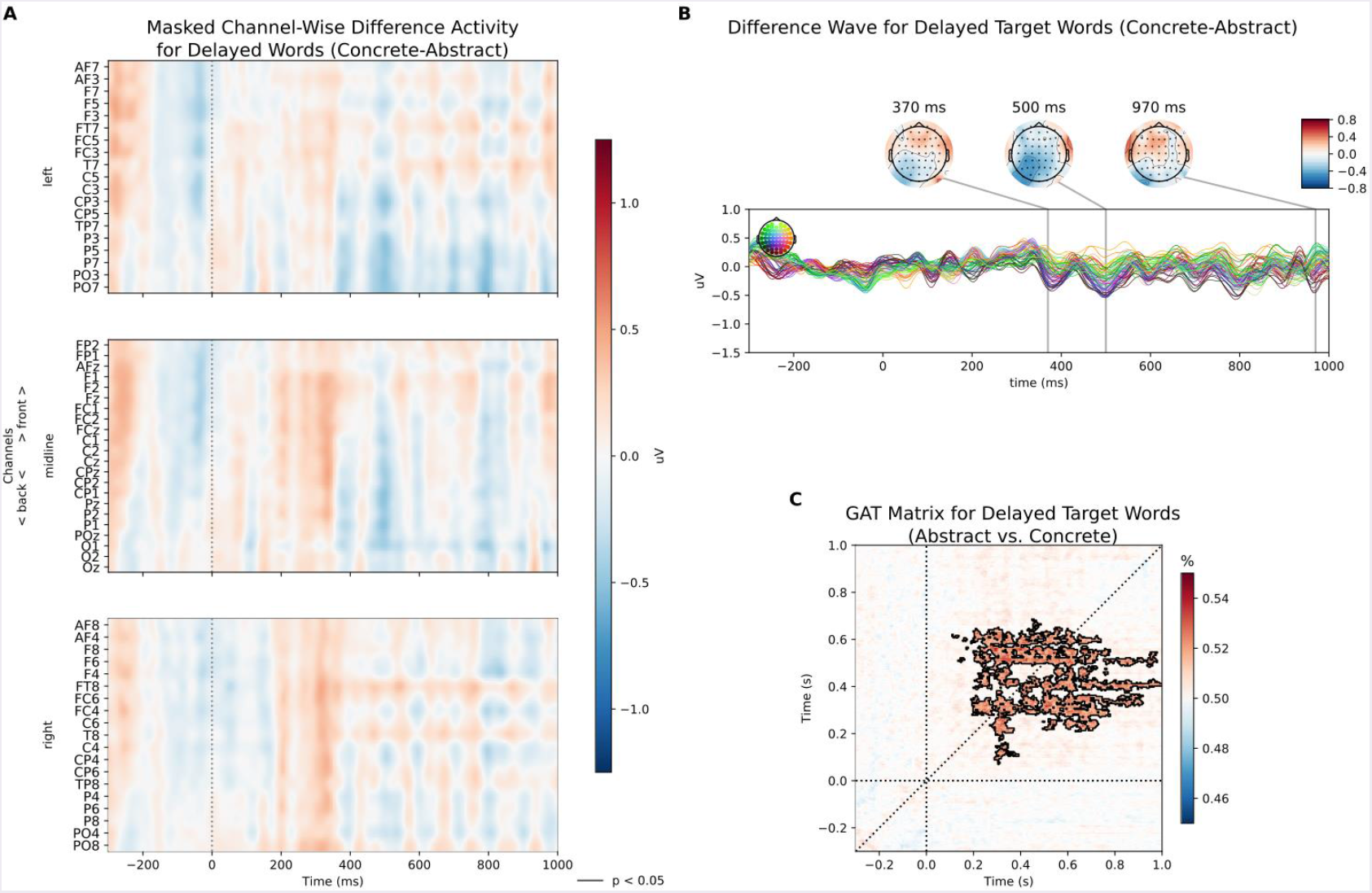
Decoding target word abstract vs. concreteness from silence (Experiment 1) Results for abstract/concrete target words, during the silent delay (1,000 ms) preceding the target word. **A)** Average channel-wise evoked potentials (concrete minus abstract) across subjects and masked for significance (p < .05), no significant differences were observed. **B)** Averaged difference wave (concrete minus abstract) across all channels. Above: for specific, representative times points, interpolated topographies demonstrate spatial structure. **C)** Generalization across time decoding with masked time points of significance (above chance; p < .05, threshold-free cluster enhancement/TFCE).

### Decoding Concrete- vs. Abstract Isolated Spoken Nouns

For the auditory word list experiment, cluster-level permutation testing revealed significant ERP differences between abstract and concrete words (p < .05). Spatially contiguous clusters were significant from 200 to 365 ms, coinciding with a negativity over right anterior electrode sites for concrete relative to abstract words (see Figure 2A & B). GAT decoding of concreteness category was also significantly above chance along the diagonal, starting at 225 ms and broadly generalizing until the end of the examined epoch (p < .05, TFCE; Figure 2C).

**Figure 2.**
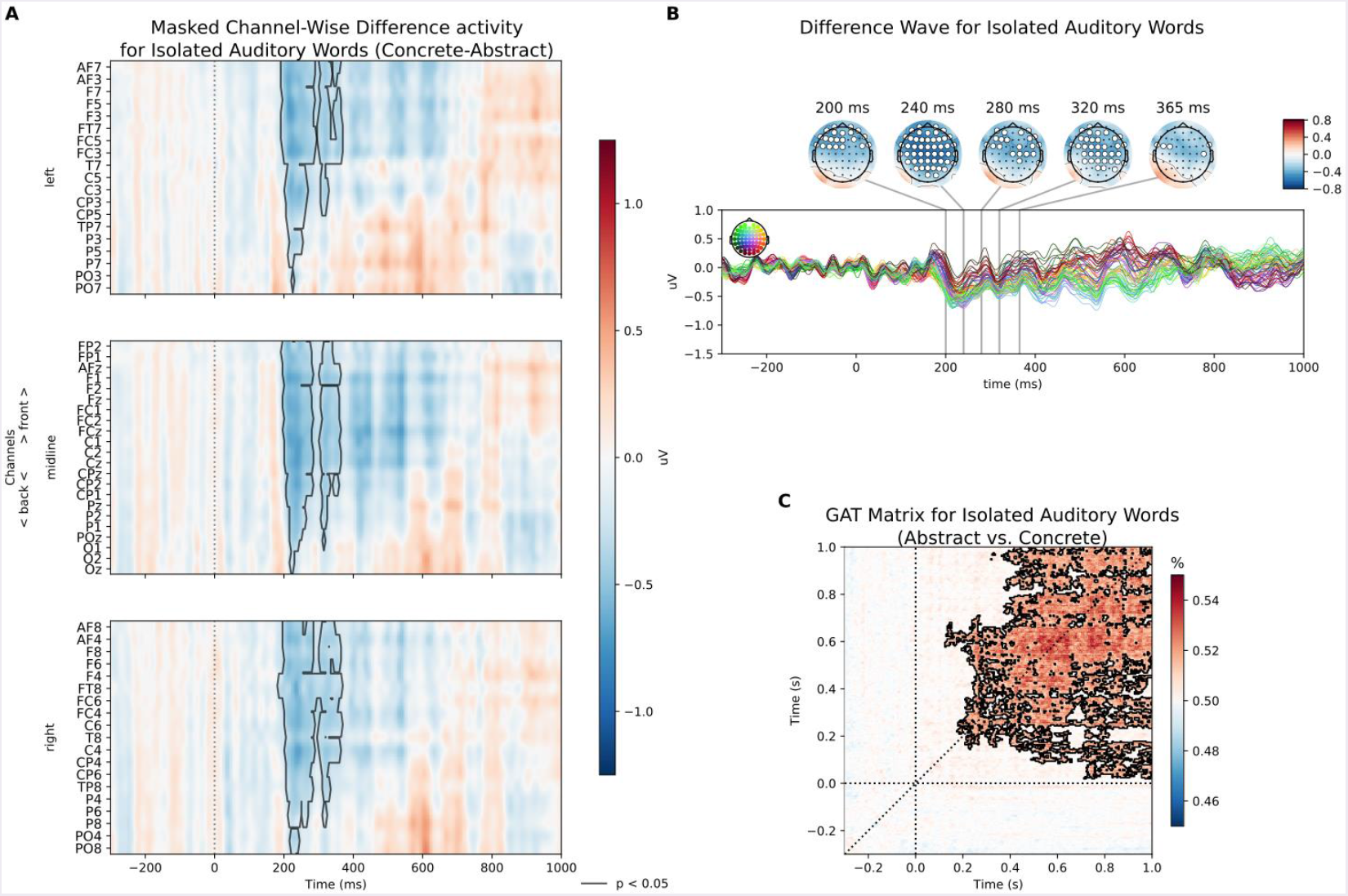
Decoding abstract vs. concreteness from isolated auditory words. (Experiment 1) Results for abstract/concrete auditory words presented in isolation. **A)** Average channel-wise evoked potentials across subjects and masked for significance (p < .05). **B)** Averaged difference wave (concrete minus abstract) across all channels. Above: for specific, representative times points, interpolated topographies demonstrate spatial structure. Channels belonging to significant clusters at the corresponding time point are highlighted in white. **C)** Generalization across time decoding with masked time points of significance (above chance; p < .05, threshold-free cluster enhancement/TFCE).

### Decoding Concrete- vs. Abstract Isolated Printed Nouns

As for auditory word presentation, ERPs elicited by visually presented words showed a broadly enhanced negativity for concrete relative to abstract words. This effect, however, started earlier and lasted longer than for auditory words, i.e., from 160 to 630 ms (see Figure 3A & B). The effect appears analogous to a typical N400 response (as is expected for a concreteness manipulation; e.g., Dufau et al., 2015). Significant GAT decoding of concreteness was observed from 250 to 955 ms (p < .05, TFCE; Figure 3C), again in a roughly square shape indicating pattern persistence.

**Figure 3.**
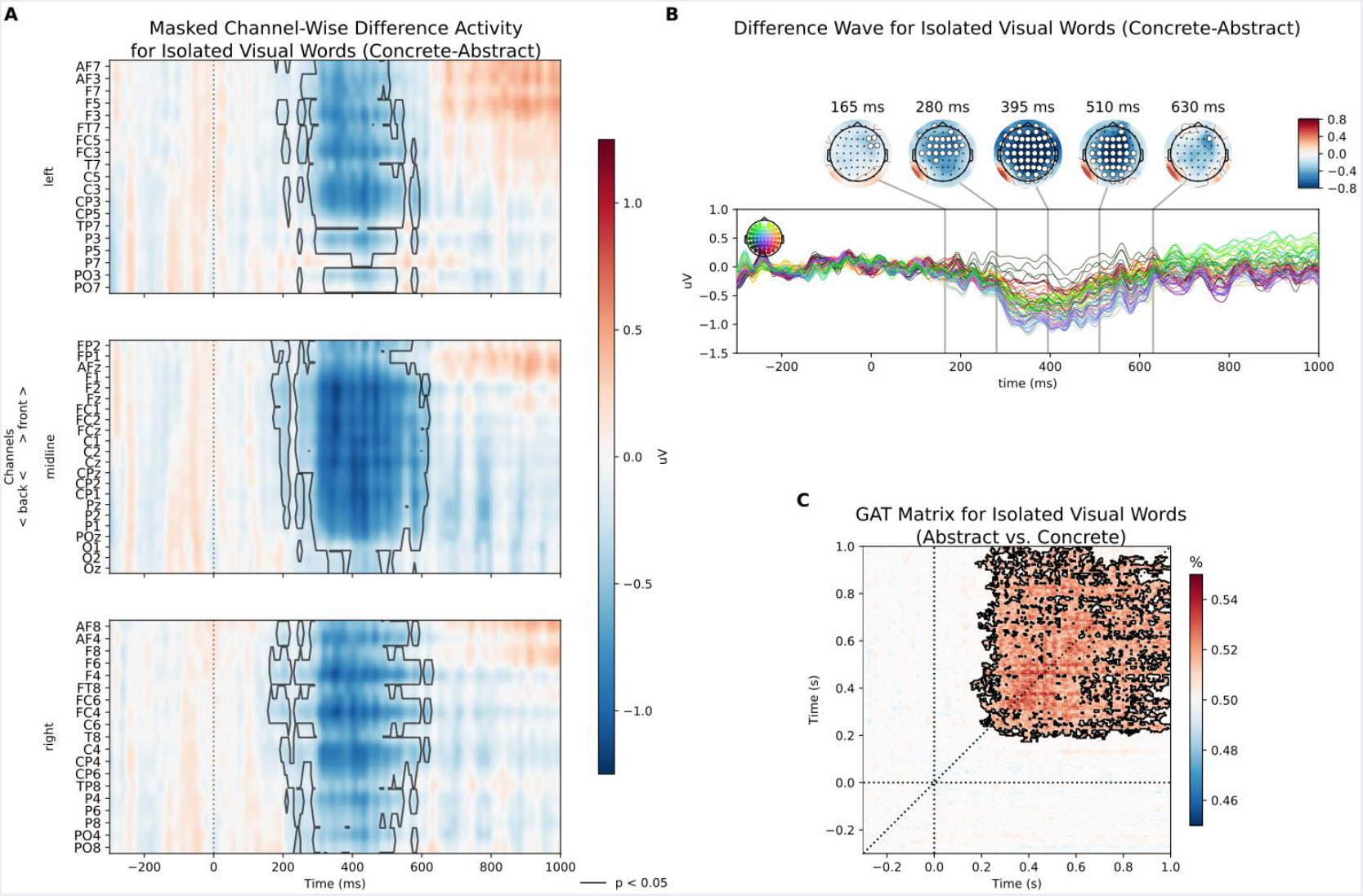
Decoding abstract vs. concreteness from isolated visual words (Experiment 1) Results for abstract/concrete visual words presented in isolation. **A)** Average channel-wise evoked potentials across subjects and masked for significance (p < .05). **B)** Averaged difference wave (concrete minus abstract) across all channels. Above: for specific, representative times points, interpolated topographies demonstrate spatial structure. Channels belonging to significant clusters at the corresponding time point are highlighted in white. **C)** Generalization across time decoding with masked time points of significance (above chance; p < .05, threshold-free cluster enhancement/TFCE).

### Between-Experiment Decoding

Finally, we tested whether the semantic predictions that – as our above-reported decoding results for the pause intervals suggest – are active prior to word presentation, share representational codes with the neural patterns elicited when words from these semantic categories are presented in isolation. This was tested using cross-decoding between experiments, i.e., training on the abstract-concrete classification in one of the single-word experiments and testing on pause-data from the sentence study. However, neither when training MVPA classifiers on the auditory, nor on the visual single word contrast could we find significant cross-decoding to EEG signals from the pause interval of the sentence experiment (TFCE, all p > .05). Classifiers trained on isolated auditory words were also not able to decode concreteness from EEG signals elicited by isolated visual words (all p > .05), and vice versa (all p > .05).

### Interim Discussion

To summarize, we found that a linear classifier could decode the categorical abstract vs. concrete nature of the expected word, from electrophysiological activity recorded during an unexpected silent period *preceding* the target word. This demonstrates the existence of pre-stimulus neural representations that code semantic expectations, providing support for the operation of predictive coding mechanisms during language processing. This finding is in agreement with a strong hypothesis derived from predictive coding theory, i.e., that the prediction error carries specific information about the expected sensory input (Friston, 2010; Clark 2013, Kok and de Lange, 2015). Under this interpretation, our results indicate that specific multivariate response patterns elicited by prediction violations in the absence of sensory input carry important information about the expected stimulus. (For discussion of potential alternative accounts, see the main Discussion section below.)

Interestingly, we found no evidence for a generalization between these omission-related semantic prediction errors and the neural representation of abstract/concreteness category elicited by words processed in isolation. This indicates that context-dependent semantic expectations are neurally computed in a different way than the semantic category membership of a word perceived without context. This is not unexpected. Isolated words are highly polysemous (compare, e.g., the many different kinds of *banks*; Hagoort, Hald, Bastiaansen, & Petersson, 2004), and sentence context quickly constrains processing paths, which is also reflected in neural signals (Federmeier et al., 2007). Thus, isolated words often activate much broader semantic information than context-embedded and disambiguated words. I.e., hearing the word “bank” in isolation activates all of the multiple meanings of bank; hearing “bank” in the context of “I took a stroll to the park and sat on a …” however allows the brain to engage a much smaller conceptual-semantic network – not only in the degree of activation, but also in the qualitative shape (i.e., in the sense of different topographies of semantic networks).

However, this line of argument is speculative and, admittedly, post-hoc. A different – albeit less likely – possibility is that some other, i.e., non-semantic aspect of the pre-omission context differed between sentence conditions and was picked up by our decoding algorithm. This type of confound would also lead to a lack of cross-experiment generalization because in that case, the signal decoded in sentences would not be a semantic signal. To exclude this possibility, a second experiment was conducted with the aim of replicating and extending the first study. Specifically, in this second study, we investigated whether a different semantic category could be read out from pre-target brain activity while controlling even better for pre-stimulus context.

## METHODS EXPERIMENT 2: DECODING WORD ANIMACY FROM SILENCE-EVOKED EEG

In this second experiment, we tested if the semantic feature ‘ animacy’ (i.e., living vs. non-living) could also be decoded from pauses preceding predictable but unexpectedly delayed words in sentences. This semantic feature was chosen as (a) it can be decoded from words presented in isolation (Chan et al, 2011), and (b) N400 results (in Polish; cf. Szewcyk and Schriefers, 2013) are in principle compatible with the predictive activation of animacy status. We thus hypothesized that the semantic category ‘ animacy’ has a specific neurophysiological representation that can be proactively elicited by a constraining sentence contexts – analogous to the conclusion we derived for concreteness based on the results of Experiment 1. As in Experiment 1, a pause of 1,000 ms was inserted prior to the target word in half of the critical sentences, and EEG measured during this silent period was the basis for the decoding analysis. Given the lack of cross-decoding between sentences and isolated words in Experiment 1 (see also the Interim Discussion, above), this second study involved only the sentence experiment.

### Participants

Forty-four native speakers of German were recruited according to the same criteria as in Experiment 1, and participated after giving informed consent according to protocols approved by the local ethics committee. Three participants were excluded because more than 33% of trials had to be rejected during artefact correction. Six participants were excluded due to faulty equipment. The final sample consisted of *n* = 35 participants (mean age = 23.20; SD = 3.39; age range 18 to 30; 8 males).

### Experimental Design: Stimuli and Procedures

Experimental procedures were identical to Experiment 1, including the synonymy judgment task. Stimuli consisted of 265 sentences, of which 108 contained a 1 s pause inserted directly preceding a target word. These sentences constrained for either an animate or an inanimate target noun (54 trials per condition; see Table 1). Sentence length varied between 4.01 and 7.98 s (mean = 5.82, SD =.90). The number of words in the sentence ranged from 6 to 17 (animate: range 6 to 17, mean = 11.07, SD = 2.68; inanimate: range 7 to 16, mean = 10.65, SD = 2.07). The delayed word occurred 49 times in the sentence final position (animate: 24, inanimate: 25), 17 in the penultimate (animate: 6, inanimate: 11), and 42 times in an earlier position (animate: 24, inanimate: 18). Word frequencies for animate words were: mean = 6.39, SD = 2.40. Inanimate: mean = 6.00, SD = 1.94. 13 synonymous sentences were created for the synonymy judgment task (not included in the analysis), and the Potsdam Sentence Corpus (PSC; 144 sentences; cf. Kliegl et al., 2004) was used as filler sentences.

Critical sentences with animate vs. inanimate target nouns were constructed and rated for target word cloze probability by 29 independent participants. Mean target noun cloze probability was .65 (SD = .29). There was no substantial difference between animate and inanimate cloze ratings (mean = .67, SD = .25 and mean = .62, SD = .32, respectively). Furthermore, the sentence context was highly constraining for the to-be-decoded semantic feature (i.e., animate vs. inanimate; mean = .93, STD = .13 vs. mean = .96, STD = .10, respectively). Finally, to minimize the potential for lingering effects of the pre-stimulus context (for the complexity of implicit matching, see Sassenhagen and Alday, 2016), sentences were created in pre-omission matched pairs across the conditions. Of course, it would not be possible to keep the pre-stimulus context exactly equal, as identical contexts would induce identical predictions. However, we constructed sentences so that the last word before the omission was identical within a pair; i.e., the same 54 words occurred in the final pre-omission position across the animate-constraining and the inanimate-constraining contexts (see Table 1). Additionally, for 22 sentences the last two words were identical, but it proved impossible to construct a sufficient number of stimuli according to this constraint.

### EEG Data Acquisition and Statistical Analysis

Experiment 2 was measured in a different laboratory, so AFz was used as ground electrode. All other aspects of data acquisition and analysis were identical to the investigation of neural activity during pauses in Experiment 1.

## RESULTS EXPERIMENT 2

ERPs elicited during pauses preceding expected animate vs. expected inanimate nouns did not differ (Figure 4A & B). However, time-generalized MVPA revealed significant decoding of animacy from 235 to 880 ms after the onset of the pause (TFCE; p < .05; see Figure 4C). Again, time-generalized decoding profiles showed substantial temporal generalization, most compatible with the sustained presence of just one discriminative pattern persisting throughout an extended time window.

**Figure 4.**
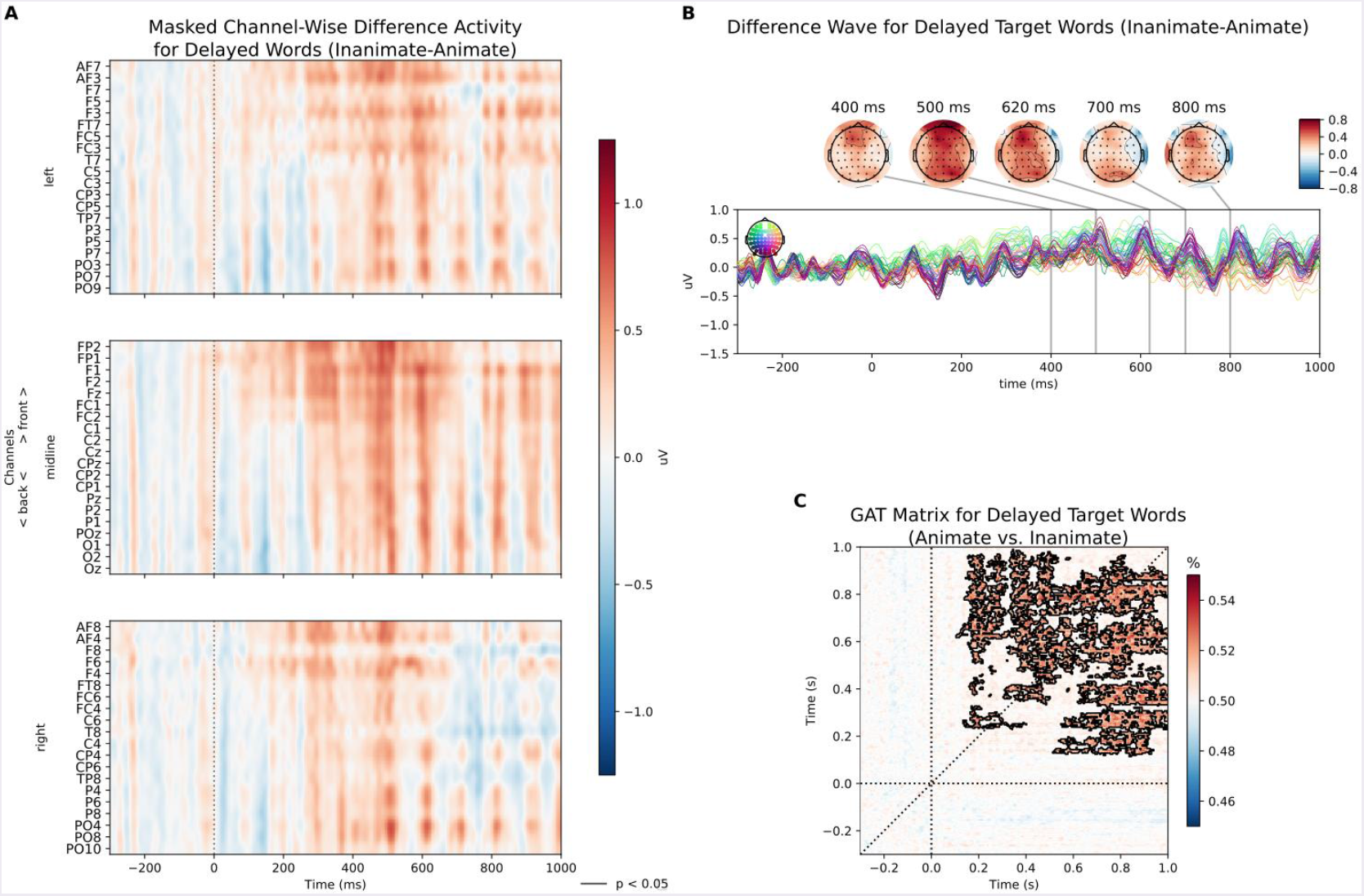
Decoding target word animacy from silence (Experiment 2) Results for animate/inanimate target words, during the silent delay (1,000 ms) preceding the target word. **A)** Average channel-wise evoked potentials (animate minus inanimate) across subjects and masked for significance (p < .05), no significant differences were observed. **B)** Averaged difference wave (animate minus inanimate) across all channels. Above: for specific, representative times points, interpolated topographies demonstrate spatial structure. **C)** Generalization across time decoding with masked time points of significance (above chance; p < .05, threshold-free cluster enhancement/TFCE).

## DISCUSSION

In this study, we adopted a predictive coding perspective on language processing and hypothesized that during sentence processing, expectations regarding upcoming linguistic input are built online, including at a semantic level. We reasoned that an unexpected delay of a word – as it frequently happens during natural language processing – is equivalent to a (temporary) omission of expected sensory input, and therefore well-suited to investigate the nature and neural realization of linguistic predictions. A strong hypothesis of the predictive coding model is that error signals elicited by prediction violations represent the difference between input and internal prediction (Friston, 2010; Clark, 2013). Thus, absence of an expected stimulus should elicit a prediction error signal that represents nothing but the internal expectation (for empirical evidence, see e.g., Kok et al., 2014; Smith and Muckli, 2010; see Kok and de Lange, 2015, for review). Applying this line of reasoning to online sentence processing, we demonstrate in two experiments that word concreteness and word animacy, respectively, could be decoded from neural activity elicited during the unexpected delay preceding an expected target word. This indicates that sentence context activates category-level semantic information of a word not yet perceived.

It is well-accepted that a constraining sentence context can support processing of subsequent words (Kutas and Federmeier, 2011). Current debates in language research are focused on whether this facilitatory effect results from predictive pre-activation of word-specific features based on higher-level (e.g., semantic) information available before the word is actually perceived, as opposed to less effortful bottom-up integration of new input (e.g., Kuperberg and Jaeger, 2015; Lau et al., 2008). Evidence for predictive pre-activation is so far mostly indirect, for example relying on prediction violation effects on determiner words prior to the supposedly predicted target noun (DeLong et al., 2005). The assumption of predictive pre-activation, however, requires that predictable linguistic information is neuronally encoded online while the sentence unfolds and before it is actually conveyed by linguistic input. Our data provide direct evidence for the predictive activation of category-level semantic expectations.

A possible alternative explanation for this finding is that our analyses may have decoded not the intended semantic feature, but some other word feature not controlled. However, having observed similar effects in two experiments on two different semantic categories greatly reduces this risk. Furthermore, even if such a confound existed, it effectively would not weaken our primary conclusion, i.e., that some aspect of an omitted word can be decoded before the word itself has been presented. A further alternative would be that above-chance decoding reflects lingering neural consequences of the preceding sentence context. In this case, decoding would not reflect internally generated linguistic predictions, but (uncontrolled) differences between pre-omission features of the sentences. This also appears unlikely, for two reasons: First, the actual words directly preceding the pause are not themselves nouns differing in animacy or concreteness. In Experiment 1, only one out of 150 pre-omission words was a noun; in Experiment 2, no nouns occurred here and the same pre-omission words were used in both conditions. Second, decoding performance is maximal starting 200-300 ms after pause onset, and not at the beginning of the delay (as would be expected for some kind of ‘ spill-over’ effect). The timing of our decoding results is more consistent with the interpretation that we decode some latent change in brain state, induced by the pre-pause sentence context, which is systematically related to the semantics of the word. This is, arguably, best described as a predictive process of some form. Even though our study, across two experiments, supports the existence of category-level semantic predictions during online sentence processing, it is necessary to follow-up this initial result and map out in more detail the nature of semantic predictions active during sentence processing. Our study demonstrates a novel approach to investigating which aspects of the preceding sentence context (reviewed, e.g., by Kuperberg and Jaeger, 2015) are causal for eliciting the semantic prediction.

While we introduced the decoded brain pattern as an omission-related prediction error, pre-word decoding of semantic categories could alternatively also reflect the neural instantiation of the linguistic prediction itself. In the absence of further evidence, our decoding results cannot unambiguously distinguish between these two alternatives (see Kok and De Lange, 2015, for a similar discussion). However, the general conclusion that can be drawn from our results, i.e., *that* semantic representations are proactively evoked during linguistic processing, is compatible with both interpretations. This supports our hypothesis that predictive coding theory – in its narrow sense – applies to language processing, including high-level linguistic phenomena like semantics and that language processing is inherently predictive. This entails that during language processing, the brain constantly builds up – predicts – complex, multi-level representations for upcoming input. Framed differently, the role of semantics in predictive processing is not simply to *enable* predictions (e.g., to guide visual search based on meaning; see, e.g., Huettig et al., 2012), but semantic features themselves are one of the units of predictive processing.

This finding dissociates semantic predictions from other well-known classes of predictive processes in the brain, such as the reward prediction error implemented by the subcortical dopamine system (e.g., Schultz et al., 1997; Holroyd and Coles, 2002), where only magnitude and sign of the prediction mismatch are encoded – suggesting that different, i.e. cortical mechanisms underlie semantic predictions. Much recent research has focused on the precise shape of the prediction error function – e.g., as Bayesian surprisal (Kuperberg and Jaeger, 2015; Frank et al., 2015) – and whether or not specific word forms are predicted based on preceding sentence context (DeLong et al., 2005; Nieuwland et al., 2017). In this context, a number of principled arguments have also been forwarded against predictive processing in language comprehension, including the potential costs of failed predictions (e.g., van Petten and Luka, 2012). Our results, however, provide very direct evidence for the online generation of category-level linguistic predictions, prior to the actual perception of the respective words. In addition, the present findings support previous proposals of proactive semantic processing (Szewcyk and Schriefers, 2013, 2018; Kwon et al., 2017) and indicate that the content of prediction error signals can be queried to address outstanding questions in language processing.

Contrary to our initial expectation, neural patterns representing semantic features of isolated words did not generalize to sentence-internal semantic predictions/prediction errors (Exp. 1). One possible reason for this failure could be a lack of statistical power for detecting small effects (Yarkoni, 2009), despite a sample size that is comparatively high for functional neuroimaging studies. A further possibility is that cross-decoding failed because semantic representations activated prior to words in sentence contexts are different from semantic representations elicited by words out of context. As discussed above (Interim Discussion), this is not entirely surprising due to, e.g., the inherent polysemy of most lexical material (Hagoort et al., 2004) and the rapidity with which it is fundamentally shaped by sentential constraint (Federmeier et al., 2007). Thus, context not only simplifies but fundamentally and qualitatively changes semantic processing.

Finally, in both experiments, the decoding effects are small in magnitude (i.e., significant but only barely above chance) and highly uneven (consider the jagged edges in Figures 1C, 2C, 3C, and 4C), prohibiting specific temporal or spatial localisation of the decoding effects. This seems to dissociate from potentially more robust pre-activation effects reported recently in semantic priming experiments (e.g., Dikker and Pylkkänen, 2013; Lau et al., 2013; Fruchter et al., 2015), but may not be surprising as we decoded from silence-evoked EEG signals in a much more naturalistic language processing context. However, this observation implicates that the observed results will be easily affected by small variations of experimental or methodological parameters. In consequence, higher-powered replication efforts are required to establish the degree of robustness of this effect. The theoretical implication of this observation is that against the background noise of the ongoing EEG, the semantic content of neural correlates of predictive processing is comparatively small.

Interestingly, we observe significant decoding based on multivariate pattern analysis in contrasts where mass univariate analysis of the ERP did not indicate any differences – neither descriptively nor in the form of statistical significance. Multivariate decoding, accordingly, showed higher sensitivity than the ERP, even though both are based on the same data. Potentially, this is because MVPA decoding identifies subject-specific patterns, thus showing more sensitivity to subtle and variable effects (King and Dehaene, 2014). For example, this situation could arise when the topography of semantic prediction error signals shows a high degree of variability between persons, so that the actual effect is not preserved during group averaging, while subject-specific decoders could still identify it.

To conclude, we provide evidence (1) for the operation of predictive coding mechanisms during on-line sentence processing, and (2) that prediction error signals do not simply indicate the magnitude of the deviance between prediction and actual input, but carry specific – in the present case: semantic – information about the expected stimulus. This strengthens predictive coding theory as a likely candidate for the principled neural mechanisms underlying online processing of language, including in the domain of (non-combinatoric, word-level) semantics.

## Acknowledgments

The research leading to these results has received funding from the European Community’ s Seventh Framework Programme (FP7/2013) under grant agreement n° 617891 awarded to CJF (ERC Consolidator Grant). The authors declare no competing financial interests.

